# Improving genetic diagnosis in Mendelian disease with transcriptome sequencing

**DOI:** 10.1101/074153

**Authors:** Beryl B Cummings, Jamie L Marshall, Taru Tukiainen, Monkol Lek, Sandra Donkervoort, A. Reghan Foley, Veronique Bolduc, Leigh Waddell, Sarah Sandaradura, Gina O’Grady, Elicia Estrella, Hemakumar M Reddy, Fengmei Zhao, Ben Weisburd, Konrad J Karczewski, Anne H O’Donnell-Luria, Daniel Birnbaum, Anna Sarkozy, Ying Hu, Hernan Gonorazky, Kristl Claeys, Himanshu Joshi, Adam Bournazos, Emily C. Oates, Roula Ghaoui, Mark Davis, Nigel Laing, Ana Topf, GTEx Consortium, Peter B Kang, Alan H Beggs, Kathryn N North, Volker Straub, James Dowling, Francesco Muntoni, Nigel F Clarke, Sandra T Cooper, Carsten G Bonnemann, Daniel G MacArthur

## Abstract

Exome and whole-genome sequencing are becoming increasingly routine approaches in Mendelian disease diagnosis. Despite their success, the current diagnostic rate for genomic analyses across a variety of rare diseases is approximately 25-50%. Here, we explore the utility of transcriptome sequencing (RNA-seq) as a complementary diagnostic tool in a cohort of 50 patients with genetically undiagnosed rare muscle disorders. We describe an integrated approach to analyze patient muscle RNA-seq, leveraging an analysis framework focused on the detection of transcript-level changes that are unique to the patient compared to over 180 control skeletal muscle samples. We demonstrate the power of RNA-seq to validate candidate splice-disrupting mutations and to identify splice-altering variants in both exonic and deep intronic regions, yielding an overall diagnosis rate of 35%. We also report the discovery of a highly recurrent *de novo* intronic mutation in *COL6A1* that results in a dominantly acting splice-gain event, disrupting the critical glycine repeat motif of the triple helical domain. We identify this pathogenic variant in a total of 27 genetically unsolved patients in an external collagen VI-like dystrophy cohort, thus explaining approximately 25% of patients clinically suggestive of collagen VI dystrophy in whom prior genetic analysis is negative. Overall, this study represents a large systematic application of transcriptome sequencing to rare disease diagnosis and highlights its utility for the detection and interpretation of variants missed by current standard diagnostic approaches.

**One Sentence Summary:** Transcriptome sequencing improves the diagnostic rate for Mendelian disease in patients for whom genetic analysis has not returned a diagnosis.

## Introduction

The advent of exome (WES) and whole genome (WGS) sequencing has greatly accelerated our capacity to identify variants that explain many Mendelian diseases in both known and new disease genes. While these technologies are mainstays in Mendelian disease diagnosis, their success rate for detecting causal variants is far from complete, ranging from 25-50% (1–4). The primary challenge of these genome-based diagnostics is that the capacity of WES and WGS to discover genetic variants substantially exceeds our ability to interpret their functional and clinical impact (5–7).

One approach to improve the interpretation of genetic variation is to integrate functional genomic information such as RNA-seq, which provides direct insight into transcriptional perturbations caused by genetic changes (8, 9). Analysis of cDNA of single genes has proven useful on a case-by-case basis to provide diagnoses to patients with Mendelian disorders (10–13), and RNA-seq has previously been used to observe the effect of pathogenic variants which were identified through DNA sequencing (14, 15). However, the use of transcriptome sequencing has not yet been assessed for discovery of pathogenic variants in a cohort of Mendelian disease patients. Such approaches have already proven useful for elucidating mechanisms of cancer and common disease (16, 17) but are not currently systematically applied to rare disease diagnosis.

Here we describe the application of this technology to the diagnosis of patients with a range of primary muscle disorders, including myopathies and muscular dystrophies, using RNA obtained from affected muscle tissue (table S1). To investigate the value of RNA-seq for diagnosis, we obtained primary muscle RNA from 63 patients with putatively monogenic muscle disorders. Thirteen of these cases had been previously diagnosed with variants expected to have an effect on transcription, such as loss-of-function or essential splice site variants, allowing us to validate the capability of RNA-seq to identify transcriptional aberrations (table S2). The remaining cohort of 50 genetically undiagnosed patients included cases for whom DNA sequencing had prioritized variants predicted to alter RNA splicing or strong candidate genes, as well as cases with no strong candidates from genetic analysis (Fig. 1A, see Materials and Methods for inclusion criteria).

**Figure 1:**
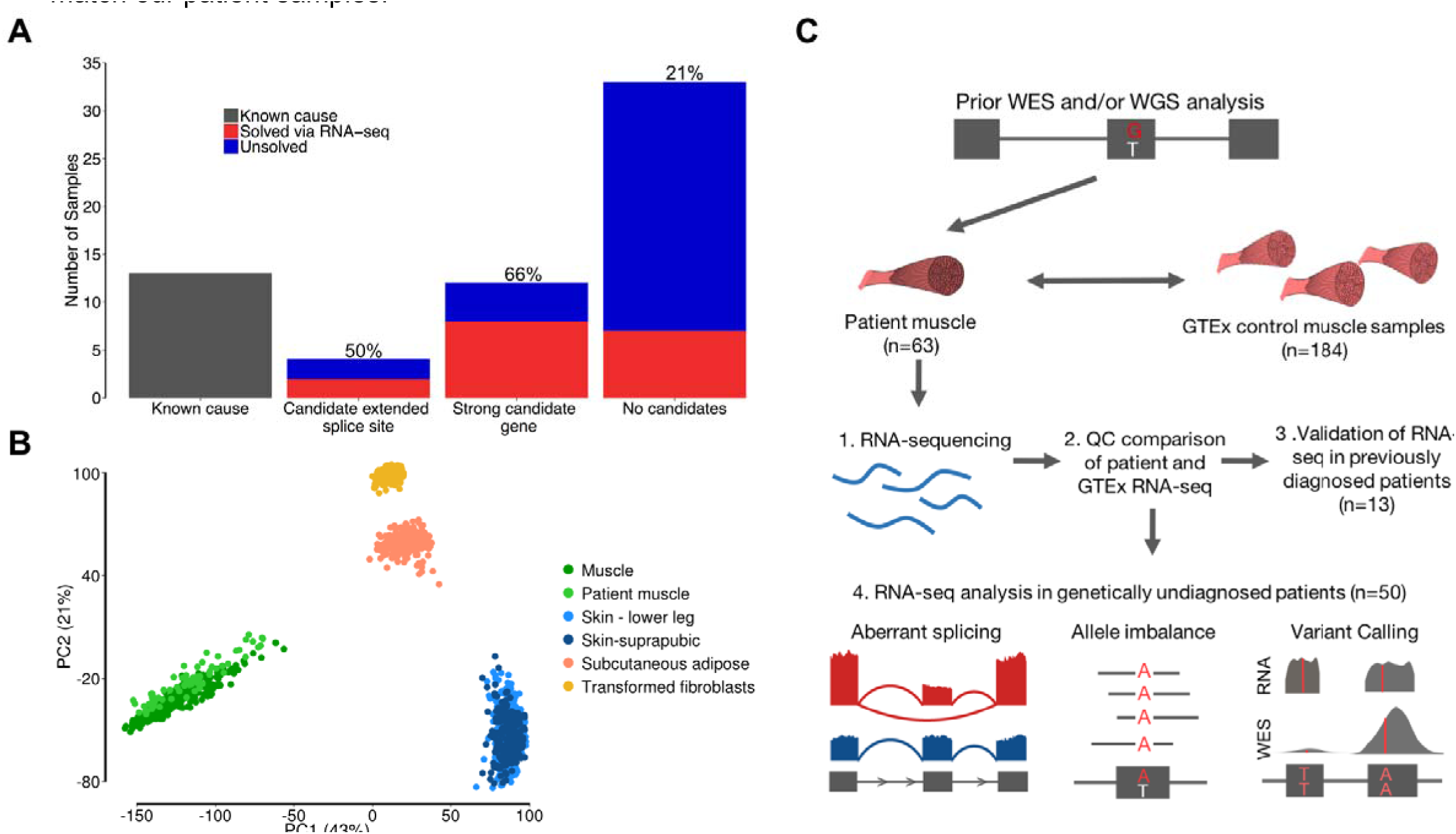
Experimental design and quality control. **A)** Overview of the number of samples that underwent RNA-seq. We performed RNA-seq oi 13 previously genetically diagnosed patients, 4 patients in whom previous genetic analysis had identified an extended splice site variant c unknown significance (VUS), 12 patients in whom genetic analysis had identified a strong candidate gene, and 34 patients with no stron; candi dates from previous analysis. RNA-seq enabled the diagnosis of 35% of patients overall, with the rate, shown above the barplots, varying depending on previous evidence from genetic analysis. **B)** PCA based on gene expression profiles of patient muscle samples passing QC (n=61) and GTEx samples of tissues that potentially contaminate muscle biopsies shows that patient samples cluster closely with GTEx skeleta muscle. **C)** Overview of experimental set up and RNA-seq analyses performed. Our framework is based on identifying transcriptione aberrations that are present in patients and missing in GTEx controls. Upon ensuring that GTEx and patient RNA-seq data were comparable we validated the capacity of RNA-seq to resolve transcriptional aberrations in previously diagnosed patients and performed analyses of aberrar splicing, allele imbalance, and variant calling in our remaining cohort of genetically undiagnosed muscle disease patients.

## Results

### Importance of sequencing the disease-relevant tissue

Recent large-scale studies have shown that gene expression and mRNA isoforms vary widely across tissues, indicating that for many diseases, sequencing the disease-relevant tissue will be valuable for the correct interpretation of genetic variation (18, 19). This is illustrated by the relative expression of known muscle disease genes in skeletal muscle, whole blood, and fibroblast samples from the Genotype Tissue Expression Consortium project (GTEx) (fig. S1) (20). The majority of the most commonly disrupted genes in muscle disease are poorly expressed in blood and fibroblasts, suggesting RNA-seq from these easily accessible tissues may be underpowered to detect relevant transcriptional aberrations in certain genes. For these reasons, we chose to pursue RNA-seq from primary muscle tissue biopsies, which are routinely performed as part of the diagnostic evaluation of undiagnosed muscle disease patients (21, 22).

### Comparison of patient RNA-seq to a muscle RNA-seq reference panel

Patient muscle samples were sequenced using the same protocol as in the GTEx project (20) and analyzed using identical pipelines to minimize technical differences, with patients sequenced at or above the same coverage as GTEx controls. From 430 skeletal muscle RNA-seq samples available through GTEx, we selected a subset of 184 samples based on RNA-seq quality metrics including RNA integrity (RIN) score and ischemic time, as well as phenotypic features such as age, BMI, and cause of death to more closely match our patient samples.

Comparison between our GTEx reference panel and patient muscle RNA-seq samples showed analogous quality metrics (table S3). Principal component analysis (PCA) of expression and splicing profiles demonstrated that patient muscle RNA-seq closely resembled control muscle when compared to tissues that potentially contaminate muscle biopsies, such as skin or fat, despite variation in the site of muscle biopsy across patients (Fig. 1B, fig. S2A, table S1). Based on this clustering, we removed two samples from analysis because their expression patterns clustered more closely with GTEx adipose tissue than muscle, consistent with tissue contamination or late-stage degenerative muscle pathology (fig. S2B). We also performed fingerprinting based on patient WES, WGS, and RNA-seq data to ensure the source of DNA sequencing and muscle RNA-seq data was the same individual.

We explored the utility of analyzing patient RNA-seq data to detect aberrant splice events and allele-specific expression and performed variant calling from RNA-seq data to identify pathogenic events or to prioritize genes for closer analysis (Fig. 1C). We also identified outlier gene expression status in patients; however, this analysis was underpowered to prioritize candidate genes in our study (fig. S3). The resulting diagnoses were made primarily through detection of aberrant splice events in patients, with information on gene-level allele imbalance playing a complementary role.

In previously diagnosed cases, manual evaluation of pathogenic essential splice site variants revealed a splice aberration such as exon skipping or extension, demonstrating that RNA-seq can help resolve the effect of variants on transcription (fig. S4A-F). To detect aberrant transcriptional events genome-wide, we developed an approach based on identifying high quality exon-exon splice junctions present in patients or groups of patients and missing in GTEx controls (code available at https://github.com/berylc/MendelianRNA-seq). We performed splice junction discovery from split-mapped reads, considering only those that were uniquely aligned and non-duplicate. To account for library size and stochastic gene expression differences between samples, we performed local normalization of read counts based on read support for overlapping annotated junctions (fig. S5A, B). We then performed filtering of splice junctions based on the number of samples in which a splice junction is observed and the number of reads and normalized value supporting that junction in each sample. Our approach successfully re-identified all known pathogenic events in patients in whom manual evaluation had revealed aberrant splicing around splice variants previously identified through genomic testing. We defined filtering parameters that selectively identified these previously known aberrant splice events and applied them to our remaining cohort of undiagnosed patients. This method resulted in the identification of a median of 5, 26, and 190 potentially pathogenic splice events per sample in ∼190 neuromuscular disease associated genes, OMIM genes, and all genes respectively (fig. S6), which required manual curation to interpret pathogenicity and led to the diagnoses made in this study.

### Diagnoses made via RNA-seq

RNA-seq allowed the diagnosis of 17 previously unsolved families, yielding an overall diagnosis rate of 35% in this challenging subset of rare disease patients for whom extensive prior analysis of DNA sequencing data had failed to return a genetic diagnosis. We also identified splice disruption in other known and putatively novel disease genes in several patients; however, due to unavailability of additional information, such as parental DNA, we could not pursue these cases further (fig. S7). Detection of aberrant splicing led to the identification of a broad class of both coding and non-coding pathogenic variants resulting in a range of splice defects such as exon skipping, exon extension, exonic and intronic splice gain, which were validated by RT-PCR analysis (Fig. 2, Table 1, Supplementary Materials and Methods). RNA-seq patterns also helped pinpoint three structural variants in *DMD* that were subsequently confirmed by WGS (fig. S8).

**Figure 2:**
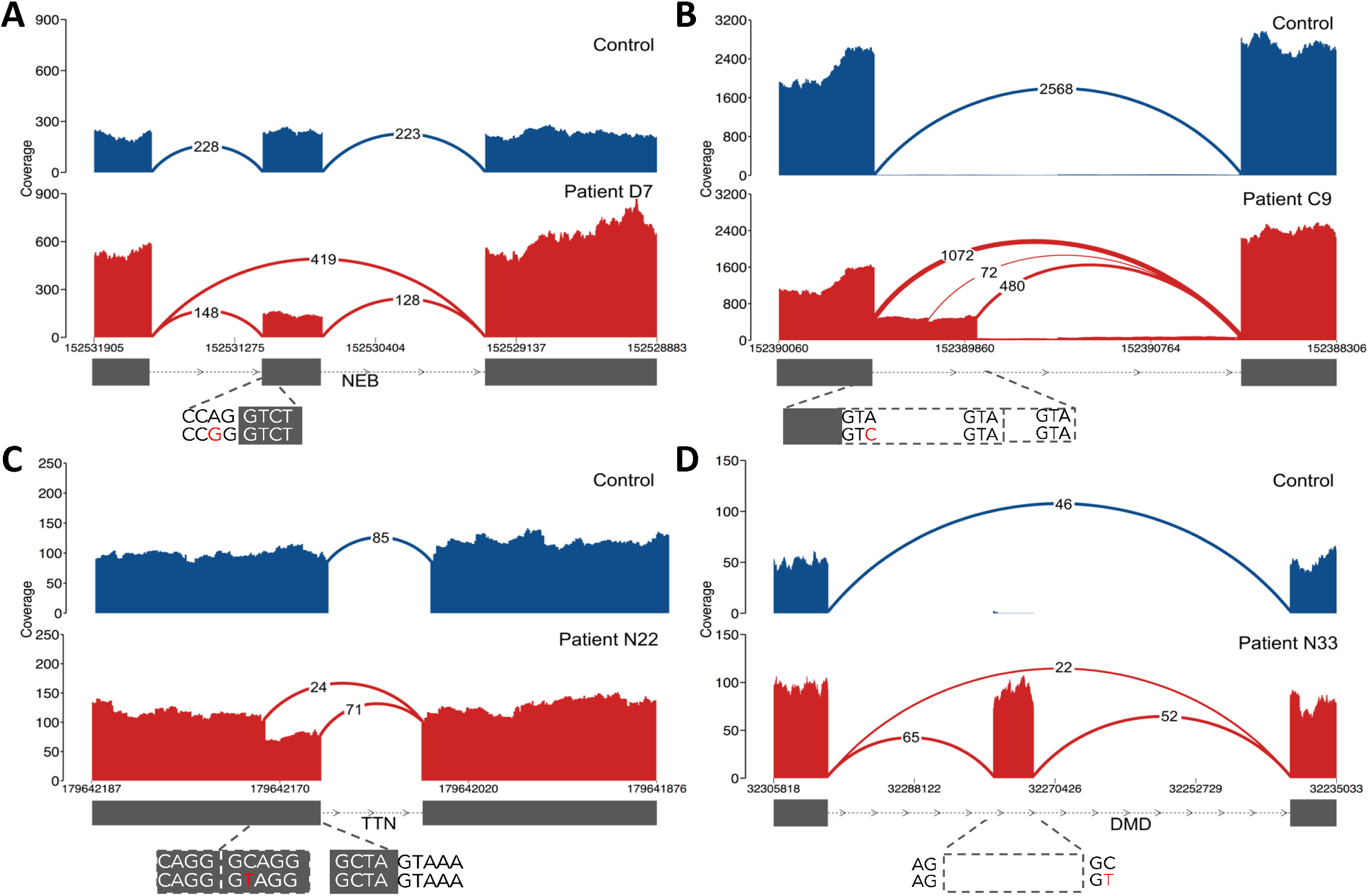
Types of pathogenic splice aberrations discovered in patients. RNA-seq identified a range of aberrations caused by both coding and non-coding variants such as **A)** exon skipping caused by an essential splice site variant in patient D7, **B)** exon extension caused by a donor +3 A>C extended splice site variant in nemaline myopathy patient C9, where disruption of splicing at the canonical splice site results in splicing from intact GTA motifs from the intron, **C)** exonic splice-gain caused by a C>T donor splice site-creating variant in patient N22 with a donor + 5G sequence context, resulting in a stronger splice motif than the existing canonical splice site, and **D)** intronic splice gain in patient N33 caused by a C>T donor splice site-creating deep intronic variant. Evidence for wild type splicing in addition to the inclusion of the pseudo exon in the patient is in line with the milder Becker’s muscular dystrophy phenotype. Splice aberrations shown in B, C, and D result in the introduction of a premature stop codon to the transcript.

**Table 1.** Diagnoses made in the study via patient muscle RNA-seq

| Patient | Phenotype | Gene | Variants | Variant Class | Effect |
| --- | --- | --- | --- | --- | --- |
| E2 | Nemaline myopathy | *NEB* | chr2:152,544,805 C>T<br>chr2:152,520,057 C>T | essential splice,<br>extended splice | exon skipping + exon<br>extension,<br>exon extension |
| C9 | Nemaline myopathy | *NEB* | chr2:152,581,432 TG>T<br>chr2:152,389,953 A>C | frameshift,<br>extended splice | exon extension |
| E4 | Fetal akinesia | *TTN* | chr2:179,586,600 CAT>C<br>chr2:179,446,219 ATACT>A | frameshift,<br>extended splice | exon skipping |
| C6 | Duchenne muscular dystrophy | *DMD* | chrX:32,366,860 A>C | intronic variant | intronic splice-gain |
| N33 | Myalgia,<br>myoglobunuria | *DMD* | chrX:32,274,692 G>A | intronic variant | intronic splice-gain |
| C7 | Becker muscular dystrophy | *DMD* | chrX:31,613,687 G>T | intronic variant | Intronic splice-gain |
| N29 | Collagen VI-related dystrophy | *COL6A1* | chr21:47,409,881 C>T | intronic variant | intronic splice-gain |
| N30 | Collagen VI-related dystrophy | *COL6A1* | chr21:47,409,881 C>T | intronic variant | intronic splice-gain |
| N31 | Collagen VI-related dystrophy | *COL6A1* | chr21:47,409,881 C>T | intronic variant | intronic splice-gain |
| N32 | Collagen VI-related dystrophy | *COL6A1* | chr21:47,409,881 C>T | intronic variant | intronic splice-gain |
| N25 | Nemaline myopathy | *NEB* | chr2:152,355,017 G>T<br>chr2:152,449,646G>A | intronic variant,<br>nonsense | intronic splice-gain |
| C11 | Congenital fiber-type disproportion | *RYR1* | chr19:38,958,362 C>T<br>chr19:38,958,372 G>A | synonymous,<br>missense | exonic splice gain |
| N22 | Multi/minicore congenital myopathy | *TTN* | chr2:179,642,185 G>A<br>chr2:179,523,240 CTTCT>C | missense,<br>frameshift | exonic splice-gain |
| C1 | Alpha dystroglycanopathy | *POMGNT1* | chr1:46,655,129 C>A<br>chr1:46,660,532 G>A | essential splice,<br>synonymous | exonic splice-gain,<br>exon skipping |
| C3 | Duchenne muscular dystrophy | *DMD* | chrX:31,790,694-31,798,498 | inversion-deletion | exon skipping |
| C2 | Duchenne muscular dystrophy | *DMD* | chrX:31,378,946-151,194,962 | inversion | splice disruption |
| C4 | Duchenne muscular dystrophy | *DMD* | chrX:32,521,820-35,180,380 | inversion | splice disruption |

Cases diagnosed in this study highlight several key advantages of RNA-seq in rare disease diagnosis to confirm the pathogenicity of variants and to detect previously unidentified variation. In four patients with previously detected extended splice site variants of unknown significance (VUS), RNA-seq confirmed splice disruption in two patients (Fig. 1A, fig. S9A, B). The variants had no observable effect on local splicingpatterns in the remaining two patients, emphasizing the value of RNA-seq in ruling out non-pathogenic VUS (fig. S9C, D).

RNA-seq also led to the identification of an additional disruptive extended splice site variant missed by exome sequencing. In a nemaline myopathy patient with one previously detected recessive frameshift variant in the *NEB* gene, RNA-seq identified an exon extension event caused by an underlying variant at the +3 position of the donor site which led to the introduction of a premature stop codon to the transcript as the second recessive allele (Fig. 2B). The exon harboring this variant was not captured in the exome kit used to screen the patient (fig. S10), underlining the utility of RNA-seq at complementing WES to identify previously undetected variants.

Synonymous and missense variants in large, variation-rich genes such as *TTN* are exceptionally challenging to interpret and are often filtered out in DNA sequencing pipelines (23, 24). With RNA-seq, we were able to assign pathogenicity to a missense variant in *TTN* and two synonymous variants in *RYR1* and *POMGNT1* (fig. S11). In patient N22, the identified missense variant created a GT donor splice site for which the consensus motif included a G nucleotide in the +5 position, known to contribute to the strength of the splice site (25, 26). The well-conserved donor +5-G motif was missing in the competing canonical splice site, thus resulting in a stronger novel splice site and gain of splicing from the exon body (Fig. 2C). A similar mechanism was observed in *RYR1*, caused by a synonymous variant in a patient carrying a second pathogenic allele in the gene (fig. S11A). In an additional patient carrying an essential splice site variant in *POMGNT1*, we identified a synonymous variant disrupting an exonic splice motif and resulting in exon skipping (fig. S11B-D).

In eight cases, RNA-seq aided in the identification of non-coding pathogenic variants. We identified splice site-creating hemizygous deep intronic variants in *DMD* that resulted in the creation of a pseudo-exon and led to a premature stop codon in the coding sequence in three patients (Fig. 2D, fig. S12). Although RNA-seq from a patient with severe Duchenne muscular dystrophy showed only splicing to the pseudo-exon (fig. S12), wildtype splicing between annotated exons was observed in two patients with a milder Becker muscular dystrophy phenotype, indicating the presence of residual functional *DMD* transcripts that explain the milder disease course. Such intronic variants are unobservable with WES and too abundant to be interpretable with WGS alone, emphasizing the utility of RNA-seq at resolving pathogenicity of these noncoding variants

In two patients with no strong candidates from WES and WGS (N22 and N25) we identified heterozygous splice disruption in two commonly disrupted recessive muscle disease genes, *NEB* and *TTN*. These genes harbor regions with highly similar sequences, the so-called triplicate repeat regions (27, 28). Due to high sequence similarity, the region has poor mapping quality, resulting in low quality variant calls that are filtered by most current diagnostic pipelines. To identify possible pathogenic variants in the triplicated regions of *NEB* and *TTN* in these two patients, we developed a method based on remapping the triplicate regions to a de-triplicated pseudo-reference and performing hexaploid variant calling (fig. S13A-C). This method was applied to available WES/WGS and RNA-seq data for all patients and identified one novel nonsense and one novel frameshift variant in *NEB* and *TTN* in these two patients, which finalized their diagnoses (N25 and N22, fig. S13D and E, respectively).

### Identification of a recurrent splice site-creating variant in collagen VI-related dystrophy

A notable example of the power of transcriptome sequencing is our discovery of a genetic subtype of severe collagen VI-related dystrophy, which is caused by mutations in one of three collagen 6 genes (*COL6A1, COL6A2*, and *COL6A3*) (21). In four patients who had previously tested negative with deletion/duplication testing and fibroblast cDNA sequencing of the collagen VI genes as well as clinical WES and WGS, we identified an intron inclusion event in *COL6A1* using RNA-seq (Fig. 3A). The splicing-in of this intronic segment, which is missing in GTEx controls and all other patients in our cohort, is caused by a donor splice site-creating GC>GT variant that pairs with a cryptic acceptor splice site 72 bp upstream, creating an inframe pseudo-exon (Fig. 3B). This variant is missing in the 1000 Genomes Project dataset (29) as well as an in-house dataset of 5,500 control WGS samples. The resulting inclusion of 24 amino acids occurs within the N-terminal triple-helical collagenous G-X-Y repeat region of the *COL6A1* gene, the disruption of which has been well-established to cause dominant-negative pathogenicity in a variety of collagen disorders (30). Of note, cDNA analysis shows that the aberrant transcript is observable in muscle but in much smaller amounts in cultured dermal fibroblasts, making the event identifiable by muscle transcriptome analysis despite being previously missed by fibroblast cDNA sequencing (Fig. 3C). Using this information, we genotyped the variant in a larger, genetically undiagnosed collagen VI-like dystrophy cohort and identified 27 additional patients carrying the intronic variant. We confirmed that the variant had occurred as an independent de novo mutation in all 16 families for whom trio DNA was available. Based on this screening, we estimate that up to a quarter of all cases clinically suggestive of collagen VI-related dystrophy but negative by exon-based sequencing are due to this recurrent de novo mutation (Supplementary Materials and Methods).

**Figure 3:**
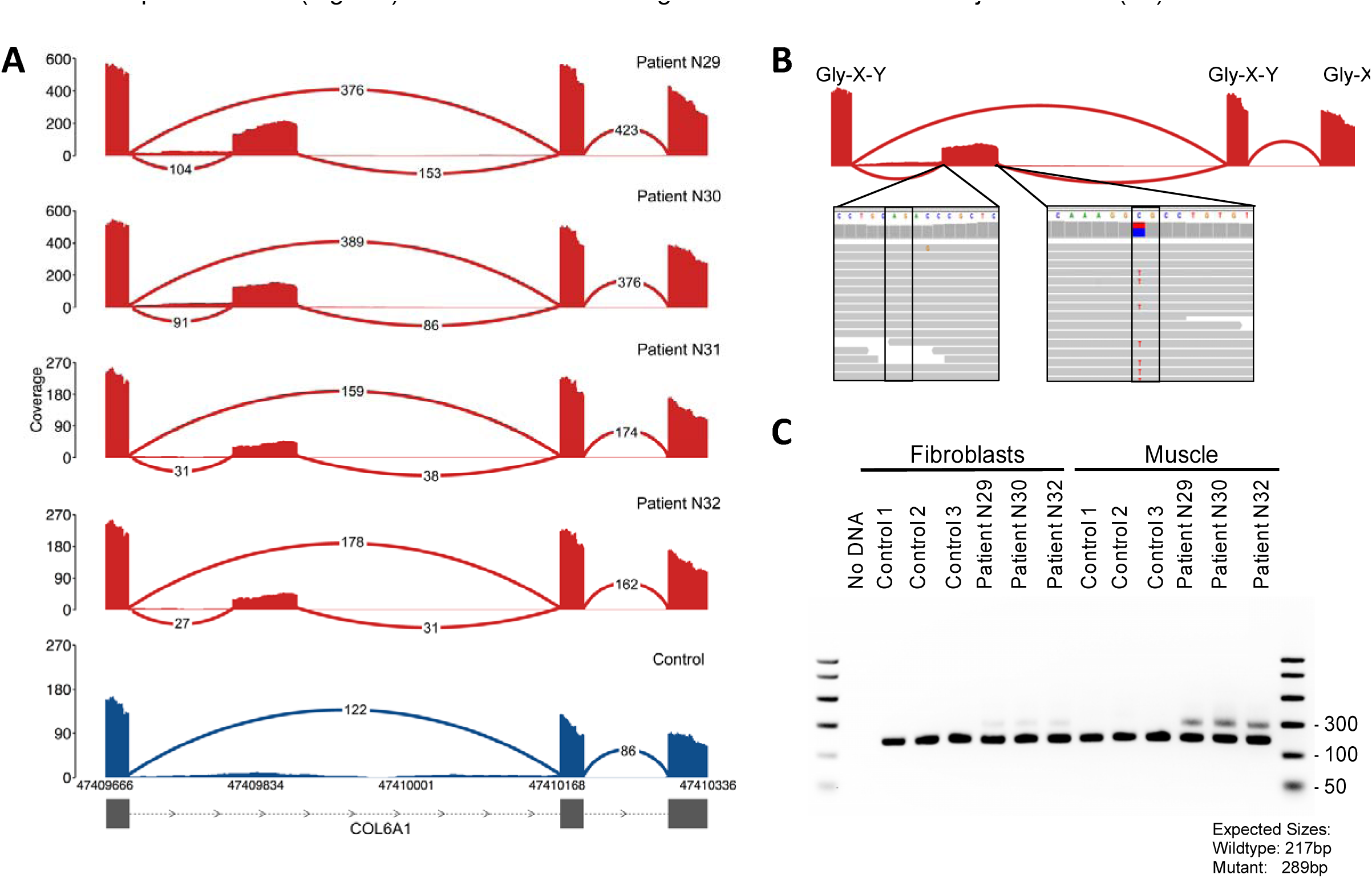
Identification of a recurrent splice site-creating variant in four collagen VI-related dystrophy patients. **A)** Splicing in of the pseudo-exo was observed in four patients in our cohort (red) and missing in all other patients and GTEx samples (blue). **B)** Inclusion of the 24 amino aci segment is caused by a C>T donor splice site-creating variant which pairs with a AG splice acceptor site 72 bp upstream. The variant is found in a CpG nucleotide context, which likely explains its recurrent *de novo* status, and disrupts the Gly-X-Y repeat motifs of *COL6A1*. **C)** The inclusion event is observable in RT-PCR amplicons from patient muscle but is found at comparatively lower levels in cultured dermal fibroblast derived from the patients, explaining why the pathogenic event was missed in all four patients through previous fibroblast cDNA sequencing.

### Evaluation of splice prediction algorithms and RNA-seq in alternative tissues

Exons harboring the pathogenic variants identified in this study show low coverage in GTEx whole blood and fibroblast samples, indicating that a majority of these diagnoses likely could not have been made using RNA-seq from these tissues (fig. S14). Furthermore, many of the diagnoses made in this study could not have been made on genotype information alone, as splice prediction algorithms alone are currently insufficient to classify variants as causal (31, 32). Although existing *in silico* algorithms correctly predicted disruption for the two extended splice site variants of unknown significance in our study, they also generated false positive predictions for the remaining two extended splice site variants with no effect on splicing (Supplementary Materials and Methods, fig. S15A). In addition, existing algorithms showed poor specificity in identifying splice site-creating coding variants, identifying on average over 100 putative splice site-creating rare variants (<1% population frequency in ExAC) exome-wide (fig. S15B).

## Discussion

Our results show that RNA-seq is valuable for the interpretation of coding as well as non-coding variants, and can provide a substantial increase in diagnosis rate in patients for whom exome or whole genome analysis has not yielded a molecular diagnosis. In our cohort, RNA-seq led to the diagnosis of 66% of patients where clinical phenotyping and DNA sequencing prioritized a strong candidate gene. In comparison, through identifying aberrant splice events found in patients and missing in GTEx controls, we were able to diagnose 21% of patients with no strong candidates from WGS or WES.

Our work illustrates the value of large multi-tissue transcriptome data sets such as GTEx to serve as a reference to facilitate the identification of extreme splicing or allele balance outlier events in patients. In the case of muscle disorders, our diagnoses were made primarily through direct identification of aberrations in splicing using the GTEx skeletal-muscle RNA-seq dataset as a reference panel. Our present work focused on identifying such aberrations in known muscle disease genes, and the considerably lower number of putatively pathogenic events identified in neuromuscular disease genes versus all genes underlines the advantage of a candidate gene list for this analysis Further improvements in filtering identified splice junctions to obtain a smaller list of candidate events will be useful to expand this work for new disease gene discovery. In addition, with increasing sample sizes and improvements in methods, RNA-seq can also be used to identify somatic variants and to detect regulatory variants upstream, through analysis of expression status and allelic imbalance.

Access to the disease-relevant tissue for many Mendelian disorders remains a major barrier for the use of transcriptome sequencing in genetic diagnosis. The RNA-seq framework developed in this study can be adapted for rare diseases where biopsies are available, such as Mendelian disorders affecting heart, kidney, liver, skin, and other tissues. For example, during the preparation of this manuscript, the application of RNA-seq to fibroblast samples for the genetic diagnosis of mitochondrial disease was reported in an unpublished preprint (33). For disorders where biopsy of the disease-relevant tissue is unattainable, analyses are possible through identification of proxy tissues using databases such as GTEx and careful consideration of the expression status of the relevant genes in the proxy tissue. Alternatively, the framework developed in this study can also enable diagnoses through reprogramming patient cells into induced pluripotent stem cells and differentiation into disease-relevant tissues of interest.

Evaluation of existing splice prediction algorithms for the splice-disrupting variants identified in the study highlights that information on DNA sequence alone does not currently match the ability of RNA-seq to identify the transcriptional consequences of variants on a genome-wide scale. The diagnoses made in our study with RNA-seq, particularly the discovery of the highly recurrent mutation in *COL6A1*, demonstrates that other such cryptic splice-affecting variants may contribute substantially to undiagnosed diseases that have evaded prior detection with exome or whole genome analysis.

Overall, this work suggests that RNA-seq is a valuable component of the diagnostic toolkit for rare diseases and can aid in the identification of new pathogenic variants in known genes as well as new mechanisms for Mendelian disease.

## Methods

### Study design

We sought to explore the utility of transcriptome sequencing as a complementary diagnostic tool to exome and whole genome analysis. We reasoned that RNA-seq would allow us to interpret variants previously identified through genetic analysis and may pinpoint genetic lesions that may have eluded DNA sequencing. To interpret transcriptional aberrations seen in patients, we obtained a reference panel of 184 sets of skeletal muscle RNA-seq data from the Genotype Expression Consortium (GTEx) project. Our framework was based on identifying transcriptional aberrations present in patients that are missing in GTEx controls. We first validated the capacity of RNA-seq to resolve transcriptional aberrations in thirteen patients with prior genetic diagnosis and then analyzed the remaining fifty genetically undiagnosed patients to detect aberrant splice events and allele-specific expression and performed variant calling from RNA-seq data to identify pathogenic events or to prioritize genes for closer analysis

### Clinical sample selection

Patient cases with available muscle biopsies were referred by clinicians from March 2013 through June 2016. Samples fell into four broad categories:

1. patients for whom previous genetic analysis had resulted in a diagnosis with at least one loss-of-function or essential splice site variant, serving as positive controls to assess the capability of RNA-seq to identify the transcriptional effect of the variants (n = 13, patient IDs starting with ‘D’).
2. patients with candidate extended splice site variants that had been categorized as variants of unknown significance for which assignment of pathogenicity would result in a complete diagnosis for the patient (n=4, patient IDs starting with ‘E’).
3. patients for whom a strong candidate gene was implicated due to either a well-defined monogenic disease phenotype, such as patients with clear Duchenne muscular dystrophy evidenced by clinical diagnosis and loss of dystrophin expression (n=6), or to the presence of one pathogenic heterozygous variant identified in a gene matching the patient’s phenotype, without a second pathogenic variant in that gene (n= 6, patient IDs starting with ‘C’).
4. patients with no strong candidates based on previous genetic analysis such as exome or whole genome sequencing (n=34, patient IDs starting with ‘N’)

Patients that fit categories 2-4 are referred to as undiagnosed prior to RNA-seq and constitute the denominator for the 35% diagnosis rate. All patients had prior analysis of exome and/or whole genome sequencing data, except two cases (patients E4 and D11) for whom targeted sequencing had identified a candidate extended and essential splice site variant, respectively. We favored cases with previous trio exome or whole genome sequencing: 29/63 patients had complete trios, with 3 additional patients having one parent sequenced. Although age of onset was not considered as an exclusion criterion, a majority of the patients in the cohort had a congenital or early-childhood onset primary muscle disorder.

Muscle biopsies or RNA were shipped frozen from clinical centers via a liquid nitrogen dry shipper and stored in liquid nitrogen cryogenic storage. Before submission to the sequencing platform, all muscle samples were visually inspected, photographed, cut into 50 μm sections on Leica CM 1950 model cryostat, and transferred to pre-chilled cryotubes in preparation for RNA extraction. When muscle arrived embedded in OCT, 8 μm transverse cryosections were mounted on positively charged Superfrost plus slides (VWR, 48311-703) and stained with hematoxylin and eosin (H&E) to assess the relative proportion of muscle versus fibrosis and adipose infiltration as well as the presence of overt freeze-thaw artifact. All samples analyzed with H&E showed muscle quality sufficient to proceed to RNA-seq.

### RNA sequencing

RNA was extracted from muscle biopsies via the miRNeasy Mini Kit from Qiagen per kit instructions. All RNA samples were measured for quantity and quality. Samples had to meet the minimum cutoff of 250 ng of RNA and RNA Quality Score (RQS) of 6 to proceed with RNA-seq library prep. A fraction of samples falling below an RQS of 6 were also submitted for sequencing. All samples submitted had a range of RQSs between 3.5-8.

Sequencing was performed at the Broad Institute Genomics Platform using the same non-strand-specific protocol with poly-A selection of mRNA (Illumina TruSeq) used in the GTEx sequencing project (20), to ensure consistency of our samples with GTEx control data. Paired end 76 bp sequencing was performed on Illumina HiSeq 2000 instruments, with sequence coverage of 50M or 100M. One sample (patient N33) was sequenced to higher depth at 500M reads to permit downsampling analysis of the effects of increasing RNA-seq depth.

### Selection of GTEx controls

GTEx data were downloaded from dbGaP (http://www.ncbi.nlm.nih.gov/gap) under accession phs000424.v6.p1. From 430 available GTEx skeletal muscle RNA-seq samples, we selected 184 samples based on RNA Integrity (RIN) score (between 6 and 9), number of non-duplicate uniquely mapped read pairs (between 35M and 75M), and ischemic time (<12 hours) to remove any samples that were outliers for these quality metrics. GTEx samples were further filtered to remove samples with known clinical conditions such as Klinefelter’s syndrome or those for whom death followed after long or intermediate term illness or medical intervention (Hardy Scale 0, 3, or 4). Overall, approximately 80% of GTEx samples with available muscle RNA-seq are above the age of 40 (median age 54) and have BMI over 25 (median BMI 27). Thus we selected samples to enrich for younger GTEx donors to more closely match our patient cohort. All samples below the age of 50 were selected, resulting in 76 samples with high quality RNA-seq data. We then added older samples back on the criterion that their BMI was below 30. This resulted in a total of 184 GTEx control samples for our reference panel, with comparable male and female sample count (105 male and 79 female). This filtering method also enriched RNA-seq data from organ donors and surgical donors as opposed to postmortem samples (72% of selected GTEx controls are derived from surgical or organ donors vs 45% in the unfiltered dataset). A full list of GTEx sample IDs used as the reference panel can be found in table S4.

### RNA sequencing alignment and quality-control

GTEx BAM files downloaded from dbGAP were realigned after conversion to FASTQ files with Picard SamToFastq. Both patient and GTEx reads were aligned using Star 2-Pass version v.2.4.2a using hg19 as the genome reference and Gencode V19 annotations. Briefly, first pass alignment was performed for novel junction discovery, and the identified junctions were filtered to exclude unannotated junctions with less than 5 uniquely mapped read support, as well as junctions found on the mitochondrial genome. These junctions were then used to create a new annotation file, and second-pass alignment was performed as recommended by the STAR manual to enable sensitive junction discovery. Duplicate reads were marked with Picard MarkDuplicates (v.1.1099).

Quality metrics for patient and GTEx RNA-seq data were obtained by running RNA-seQC (v1.1.8) on STAR aligned BAMs (34). PCA on gene expression was performed based on RPKM values calculated by RNA-seQC. Two samples (D6 and N3) were removed due to outlier status in PCA, consistent with a high proportion of non-muscle tissue in the samples (fig. S2B). For GTEx samples, the expression and exon-level read count data were downloaded from dbGAP under accession phs000424.v6. For PCA of exon inclusion metrics, we obtained PSI values for GTEx samples as described in (35).

To ensure that patient DNA and RNA data were identity-matched, we compared variants identified in WES, WGS, and RNA-seq data. WES, WGS, and RNA-seq data were joint-genotyped for a set of ∼5,800 common SNPs collated by Purcell et al. (36) using Genome Analysis Toolkit (GATK) HaplotypeCaller package version 3.4. We then calculated pairwise inheritance by descent (IBD) estimates between DNA and RNA-seq data using PLINK (v1.08p). Relatedness coefficients for WES, WGS, and RNA-seq data from the same individual ranged from 0.67-1.00 across our samples (mean = 0.9), compared to a range of 0-0.18 (mean= 0.001) for non-matching individuals, confirming that the sources for DNA and RNA-seq were the same for each patient in our dataset.

### Exome and whole genome sequencing

Whole exome sequencing on DNA samples (>250 ng of DNA, at >2 ng/μl) was performed using Illumina or Agilent SureSelect v2 exome capture. The exome sequencing pipeline included sample plating, library preparation (2-plexing of samples per hybridization), hybrid capture, sequencing (76 bp paired reads) and sample identification QC check. Hybrid selection libraries covered >80% of targets at 20x with a mean target coverage of >80x. The exome sequencing data were de-multiplexed, and each sample’s sequence data were aggregated into a single Picard BAM file. Whole genome sequencing was performed on 500 ng to 1.5 μg of genomic DNA using a PCR-free protocol. These libraries were sequenced on the Illumina HiSeq X10 with 151 bp paired-end reads and a target mean coverage of >30x.

Exome and genome sequencing data were processed through a Picard-based pipeline, using base quality score recalibration (BQSR) and local realignment at known indels. The BWA aligner was used for mapping reads to the human genome build 37 (hg19). Single Nucleotide Polymorphisms (SNPs) and insertions/deletions (indels) were jointly called across all samples using GATK HaplotypeCaller. Default filters were applied to SNP and indel calls using the GATK Variant Quality Score Recalibration (VQSR), and variants were annotated using Variant Effect Predictor (VEP v78); additional information on this pipeline is provided in Supplementary Section 1 of (37). The variant call set was uploaded to the seqr analysis platform (seqr.broadinstitute.org) to perform variant filtering using inheritance patterns, functional annotation, and variant frequency in reference databases including ExAC (37) and 1000 Genomes (29).

### Identification of pathogenic splice events

Splice junctions were identified from split-mapped reads, considering only uniquely aligned, non-duplicate reads that passed platform/vendor quality controls. For each splice junction we noted:

1. the genomic coordinates
2. the gene in which the junction was observed based on Gencode v.19
3. the number of samples in which the splice junction was observed
4. the number of total reads supporting the junction in 245 samples (184 GTEx and 61 patient)
5. the per-sample read support for the junction

We then performed local normalization of per-sample read support based on the support for the highest shared annotated junction (fig. S5A). For example, an exon-skipping event harbors two annotated exon-intron junctions, and we normalize this by the maximum of read count support for canonical splicing at these two wildtype junctions. This local normalization allows for filtering low-level mapping noise and accounts for stochastic gene expression and library size differences between samples (fig. S5B).

To identify pathogenic splice events, splice junctions in protein coding genes were filtered in terms of the number of samples a splice junction is present in and the number of reads and the normalized value supporting that junction. Specifically, we defined a sensitive cutoff at which an aberrant splice event is seen with at least 5% of the read support compared to the shared annotated junction, with at least 2 reads supporting the event. We also required a splice junction to contain at least one annotated exon-exon junction, indicating that the event was spliced into an existing transcript (fig. S5A). We performed analysis on a per-sample basis, each time requiring the normalized value of a given splice junction to be maximum in that sample and twice that of the next highest sample, allowing us to search for unique events in the patient.

All candidate pathogenic splice events were manually evaluated using the Integrative Genome Viewer (IGV). This resulted in the identification of aberrant splicing at 8/9 pathogenic essential splice site variants and resulted in the diagnosis of 10/17 patients in the study. A splice aberration was not observed around an essential splice site variant found in *TTN* in patient D5 due to insufficient number of reads mapping to the local region (fig. S4E). We extended filtering parameters to identify splice junctions present in fewer than 10 samples, but with high read support in each sample, allowing us to identify the intronic splice-gain event present in 4 patients in *COL6A1* (Fig. 3A). We note that this approach would also identify putatively pathogenic splice aberrations for which there are GTEx carriers. The remaining 3 Duchenne muscular dystrophy patients were diagnosed through manual analysis of splicing patterns in *DMD* and resulted in the identification of splice disruption. Overlapping structural variants at these regions were confirmed by subsequent WGS (fig. S8).

### Statistical analysis and code availability

Our approach for evaluating outlier status for allele imbalance in patients involved defining the 95% confidence interval (95% CI: mean ± 2 SD) of mean allele balance in GTEx individuals for each gene and identifying patients for whom the gene-level allele balance fell outside of the range. Comparison between GTEx and patient RNA-seq data quality metrics relied on a t-test for significance. Data processing, analysis, and figure generation were performed using scripts written in Python 2.7 and R 3.2; code for identifying and filtering splice junctions and for variant calling in the triplicate regions of NEB and TTN is available at https://github.com/berylc/MendelianRNA-seq.

## Supplementary Materials and Methods

Fig S1: Expression of commonly disrupted muscle disease genes in muscle, blood, and fibroblasts

Fig S2: PCA based on PSI metrics and gene expression of GTEx and patient samples

Fig S3: Overview of results from expression outlier analysis

Fig S4: Evaluation of RNA-seq around pathogenic essential splice site variants previously identified by genetic analysis

Fig S5: Overview of splice junction filtering approach

Fig S6: Number of potentially pathogenic splice events identified per patient

Fig S7: Examples of splice disruption in patients with no diagnosis at the completion of the study

Fig S8: Identification of aberrant splicing overlapping structural variants with RNA-seq

Fig S9: Resolving the effect of extended splice site variants with RNA-seq

Fig S10: Identification of a splice-disrupting extended splice variant missed by prior WES

Fig S11: Assignment of pathogenicity to missense and synonymous variants with RNA-seq

Fig S12: Identification of pathogenic noncoding varants with RNA-seq

Fig S13: Overview of triplicate region remapping

Fig S14: Comparison of the number of reads aligning to exons harboring pathogenic variants identified in the study in GTEx muscle, whole blood, and fibroblast tissues

Fig S15: Evaluation of splice prediction algorithms

Fig S16: Identification of allele imbalance with RNA-seq

Table S1: Overview of clinical cases that underwent RNA-seq (provided as an Excel file)

Table S2: Summary of patients previously diagnosed by genetic analysis

Table S3: Comparison of quality metrics between patient and GTEx RNA-sequencing samples

Table S4: List of GTEx control skeletal muscle RNA-seq samples (provided as an Excel file)

Table S5: PCR conditions and primers used for RT-PCR validation of splice aberrations identified via RNA-seq and Sanger sequencing of cDNA

Table S7: PCR conditions and primers used for genomic Sanger sequence validation of variants identified in patients

## Acknowledgements

Sequencing and analysis was provided by the Broad Institute of MIT and Harvard Center for Mendelian Genomics (Broad CMG). We thank Hayley Brooks, Danielle Sookiasian Meganne E. Leach, Daniel Ezzo, Jahannaz Dastgir, Anne Rutkowski, Carla Grosmann, Chamindra Konermans, Sophia Ceulemans, Mary-Lynn Chu, Ellen Moran and Katherine Matthews for sample collection and quality control. We thank Carrie Miceli, Stan Nelson Victor Rusu and David Altshuler for sharing of control cell lines and plasmids.

## Author contributions

B.B.C, T.T., D.G.M conceived and designed the experiments. B.B.C and T.T analyzed RNA-seq data. J.L.M, Y.H, A.Bo, and M.D. designed and performed validation experiments. B.B.C, M.L, S.D, A.R.F, L.W, S.S, G.O’G, H.M.R, E.O, R.G, S.T.C and C.G.B. analyzed exome and whole-genome data. S.D, A.R.F, V.B, L.W, S.S, G.O’G, E.E, H.M.R, A.S, H.G, K.G.C, E.O, R.G, N.G.L, A.T, A.Be, P.B.K, K.N.N, V.S, J.D, F.M, N.F.C, S.T.C. and C.G.B. provided patient samples and clinical information. The CMG and GTEx provided sequencing support for patient and control DNA and RNA sequencing. F.Z, B.W, K.J.K, A.O’D-L, D.B. and H.J. contributed reagents/materials/analysis tools. J.L.M, T.T, M.L, S.D, A.R.F, V.B, L.W, S.S, K.J.K, A-O’D-L, E.O, N.G.L, A.T, J.D, C.G.B and S.T.C. critically evaluated the manuscript. B.B.C. and D.G.M. wrote the manuscript.

## Materials and Data Availability

Patient sequencing data generated as part of this study were deposited in dbGAP under accession ID phs000655.v3.p1. GTEx transcriptome sequencing data can be obtained from dbGAP under accession ID phs000424.v6.p1. Code for splice junction discovery, normalization, and filtering is available on https://github.com/berylc/MendelianRNA-seq. List of OMIM and neuromuscular disease genes used for splice detection and ASE analysis can be found at https://github.com/macarthur-lab/omim and https://github.com/berylc/MendelianRNA-seq, respectively.

